# Efficient incorporation of dendrites into a large-scale cortical model reveals their surprising role in sharpening optogenetic responses

**DOI:** 10.1101/2025.08.29.673020

**Authors:** David Berling, Tibor Rózsa, Willem Wybo, Ján Antolík

## Abstract

Single-photon optogenetics enables chronic wide-field stimulation of cortex, facilitating large-scale manipulation of neural code to study cortical processing and advance neuroprosthetics. However, access to neural codes organized at fine spatial scales is compromised by the horizontal spread of stimulation-evoked cortical activity. Overcoming this limitation requires a quantitative understanding of the mechanisms contributing to spread, which include light scattering, dendritic activation, and synaptic transmission. We addressed this with morphology-aware simulations of optogenetic stimulation in a functionally-detailed network model of primary visual cortex. We find that synaptic transmission extends activation by 37–50% beyond the illuminated area, while, paradoxically, neuronal morphology sharpens activation, as apical dendrites sample from the superficial cortex, which is less affected by light dispersion. This unexpected sharpening enhances the fidelity of stimulation with spatially distributed patterns. Our study offers guidance for optogenetic interventions targeting topographically organized neural codes and provides a computational testbed to interpret such experiments.

## Main

Wide-field implantable light stimulators now enable long-term optogenetic manipulation of neuronal circuits across large cortical volumes [1, 2]. This opens new possibilities for the chronic investigation of brain development and of sensory and motor code in meaningful cortical volumes [3, 4, 5, 6], as well as for the design of brain interfaces for sensory restoration [7, 8]. With stimulation precise enough to target individual cortical columns, this technology may harness topographically organized functional representations, as orientation or position in vision, to elicit controlled, complex sensory percepts [9, 10, 11, 12, 13]. In practice, implementing advanced, functionally specific stimulation strategies proves challenging, as optogenetically evoked activity spreads beyond the targeted cortical volume, hindering the selective activation of individual cortical columns [3, 7, 4]. The activity spread is not only driven by the limited spatial precision of light delivery itself, but also by dendritic activation and synaptic network transmission, as illustrated in Fig. 1. To formulate effective stimulation protocols and ultimately refine the precision of optogenetic interfaces, the relative contributions of the aforementioned factors must be identified and their mechanisms understood. In this study, we use large-scale spiking network simulations to disentangle these sources, and determine their impact on functionally specific stimulation in primary visual cortex.

**Fig 1.**
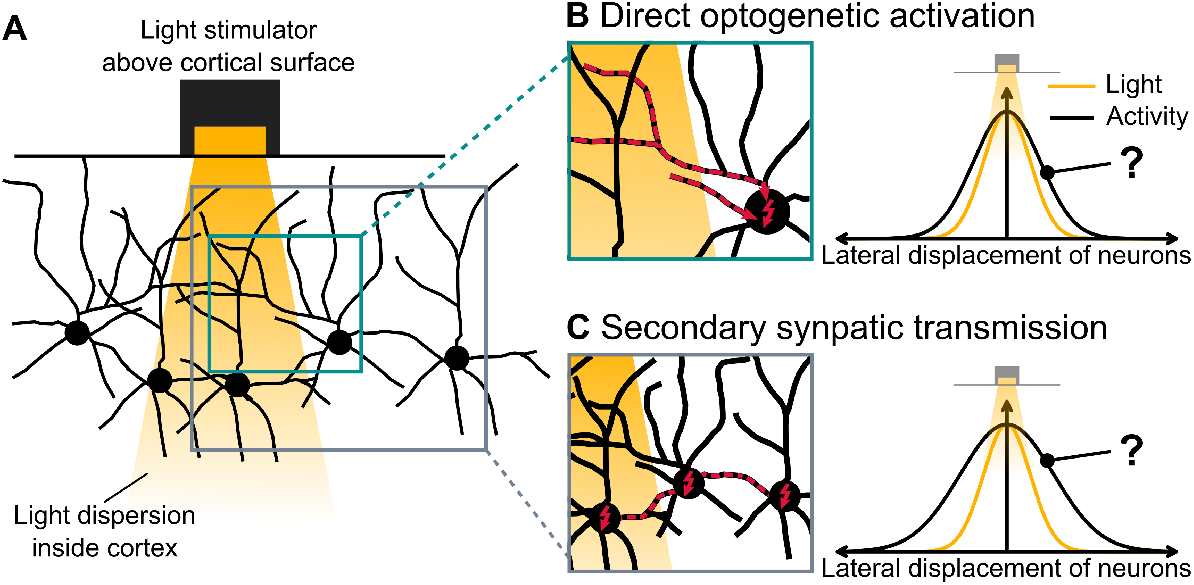
Mechanisms driving horizontal propagation of optogenetic activation inside the cortex. **(A)** Due to scattering, light introduced at the cortical surface broadens with depth, activating neurons beyond the at the surface illuminated area. Highest light intensity within a given cortical layer is observed centrally below the stimulator and decays with lateral distance. **(B)** Neurons can be directly activated via opsin-mediated conductances in their membrane when their soma or dendrites are exposed to light. Because dendrites often extend laterally, this direct activation can reach neurons outside the light beam. **(C)** In addition, activation propagates further through synaptic connections between neurons, expanding the affected area even more. The relative contributions of direct optogenetic activation and synaptic transmission to horizontal activity propagation remain unclear.

Previous computational studies have characterized factors contributing to activity spread individually, showing that neuronal morphology impacts activation in a complex, layer-dependent manner [14, 15, 13], while synaptic transmission can have both facilitatory [16] and suppressive effects [12]. However, the activation patterns caused by neuron morphology interact with the spatially organized network connectivity of the cortex, influencing how evoked activity spreads via synaptic transmission. To capture this interaction, we developed an integrated model that accounts for all three key factors: light dispersion, neuron morphology and network activity dynamics. To do so, we first overcame the bottleneck of simulating detailed neuron morphology in large-scale network models by developing an efficient approximation of morphology effects involved in optogenetic stimulation. This approximation condenses spatially distributed optogenetic input and dendritic filtering into a single somatic conductance. Combined with a point-neuron model capturing voltage dynamics, it enables efficient morphology-aware simulations of optogenetic stimulation in large cortical networks.

We then applied this model to simulate optogenetic stimulation of layer 2/3 pyramidal neurons in our state-of-the-art model of columnar primary visual cortex [17]. Our results suggest that paradoxically, morphology sharpens optogenetically evoked activity by about 17-24 % due to the presence of apical dendrites close to the cortical surface. In contrast, network transmission widened the radius of neurons activated relative to the ones receiving direct optogenetic input by about 37-50 %. This widening was more pronounced for narrower light inputs than for broader ones. Nevertheless narrow inputs remained more effective in spatially separating the evoked responses to two nearby local light stimulations, suggesting an advantage of narrow light input for inducing complex cortical activation patterns.

Our study equips the computational research community with a broadly applicable and efficient method for integrating morphology-aware optogenetic stimulation into large-scale brain models. Using these tools, we assessed key mechanisms preventing spatially precise activation of cortex, finding that synaptic transmission and light dispersion drive, while neuronal morphology suppresses horizontal activity spread. Overall, we demonstrate how large-scale detailed computational modeling can advance the understanding of brain interventions and guide the development of neurotechnologies for sensory rehabilitation and brain-computer interfaces.

## Results

This study proceeded in two steps. First, we derived and validated an efficient approximation of optogenetically driven input to morphologically-detailed cortical model neurons. Second, we integrated this approximation into an existing large-scale point-neuron model of the primary visual cortex and investigated how neuron morphology and synaptic network transmission impact the horizontal spread of optogenetically evoked activity under varying stimulation conditions.

To reduce the high computational demand of detailed optogenetic stimulation simulations in the initial step, we targeted two key aspects. First, we decreased the spatial resolution at which light intensity, opsin-induced conductance, and the resulting voltage dynamics are computed along the neuronal morphology. Second, we approximated dendritic filtering of the optogenetic input as an effective somatic conductance, thereby bypassing the need to simulate full dendritic integration.

### Reduced spatial sampling of light intensity and optogenetic conductance

Optogenetic responses strongly depend on the relative position of the light source to the neuron, reflecting the neuron’s inhomogeneous morphology (Fig. 2 A). To preserve these morphological dependencies while reducing spatial sampling to save computation time, the selected points at which the light intensity is sampled must evenly cover the entire morphology. We achieved this by choosing locations on the morphology that were closest to nodes of a Cartesian grid. Using a grid length spacing of 50 *µ*m, we reduced the number of sampling points from 1820 to 118, while maintaining a well-distributed representation of the neuronal structure (Fig. 2 B).

**Fig 2.**
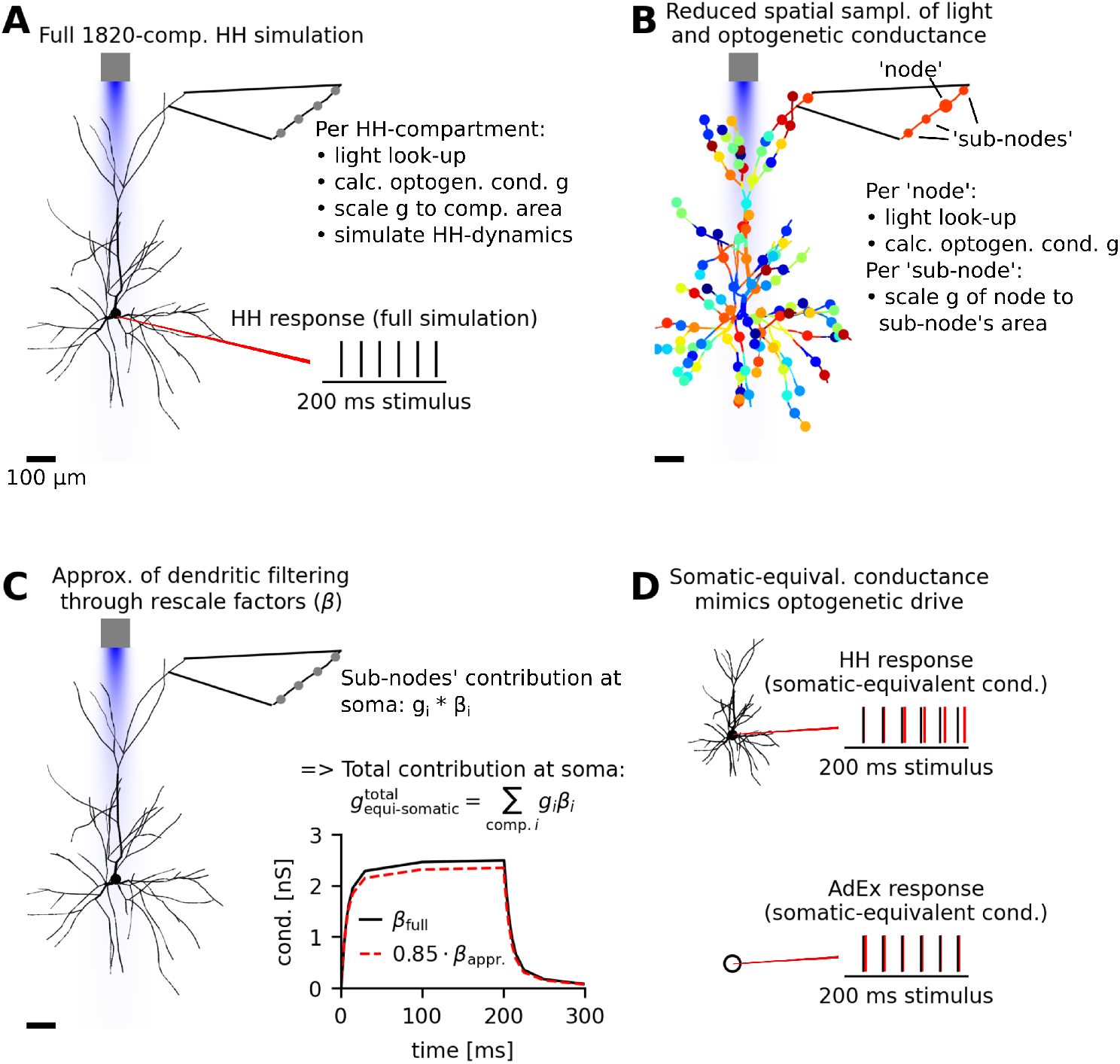
Efficient approximation of neuron-morphology effects in optogenetic responses. **(A)** Simulation of optogenetic stimulation of a layer 2/3 pyramidal cell using a 1820-compartment Hodgkin-Huxley(HH)-type model, which serves as reference for the reduction. Simulation involves light stimulus intensity read-out at each compartment and its conversion into optogenetic conductance via a Markov-state model for ChrimsonR. Voltage dynamics are simulated in all compartments to obtain the spiking response at the soma, shown in the lower right inset. **(B)** The neuron morphology of the reference layer 2/3 cell is spatially reduced from 1820 to 118 locations represented by large colored dots, which we will refer to as ‘nodes’. At the nodes, the light intensity is read out and converted into optogenetic conductance of a single ChrimsonR channel *g*. Subsequently, the single channel conductance of the nearest node is used to calculate the optogenetic conductance of all nodes and ‘sub-nodes’ (small dots) using their membrane area and optogenetic expression level. Jointly, nodes and sub-nodes corresponds to the entire set of reference model compartments. **(C)** Dendritic filtering is approximated by rescaling the distributed optogenetic inputs with rescale factors *β*. Summation over all compartments yields the somatic-equivalent conductance, which represents the aggregate effect of the optogenetic stimulation at soma. The rescale factors *β*_*i*_ depend on the coupling between the input compartment and all other locations in the morphology. To reduce computation time, we here neglected all couplings beyond the direct one between the input compartment and the soma. This approximation causes a uniform overestimation of the rescale factors over time, which we compensated through a 0.85 scale factor, as shown in the inset. **(D)** Simulation of the somatic-equivalent conductance from (C) as the only input to the reference pyramidal neuron results in a matching spike response (red) to the full simulation (black) from (A). Similarly, simulation of the somatic-equivalent conductance as the only input to an adaptive exponential integrate-and-fire (AdEx) point-neuron model results in a matching spiking response (red) to the full simulation response (black). Dynamics of the AdEx neuron model were fitted to the reference model beforehand. Optogenetic stimulation was simulated as 200 ms constant illumination with an optical fiber model (diameter (*d*) = 50 *µ*m diameter, numerical aperture (*NA*) = 0.1) at 0.2 mW/mm^2^ output light intensity.

### Approximating optogenetic conductance in the entire morphology as single conductance at the neuron’s soma

To avoid simulations of voltage dynamics in many compartments representing a single neuron, we approximated dendritic filtering of the optogenetic input to represent it as single somatic input conductance with an approach inspired by Wybo et al. [18], see Fig. 2 C. To analytically evaluate coupling between input locations and the soma in the initial HH-type model, we replaced all active conductances at rest with passive leak conductances and adjusted their reversal potentials to preserve resting voltages across the morphology. This yielded an approximate passive representation of the neuron’s response dynamics. Based on the electrical coupling of dendritic sites to soma we derived rescale factors (see *Methods*), which enabled rescaling any optogenetic input received along the morphology to represent its effect at soma. We summed inputs across the entire morphology to compute the total effect of distributed optogenetic conductance at the soma, which we termed *somatic-equivalent conductance*. By neglecting coupling between input locations, we avoid the computational cost of inverting an *N* × *N* matrix, where *N* is the number of input locations/compartments. The resulting underestimation of rescale factors due to omitted inter-compartment interactions can be corrected using a global factor, *α* = 0.85, determined through a comparison to full compartmental interaction calculations, see Fig. 2 C, bottom right.

Overall, the reduction enabled a 26-times faster computation of the layer 2/3 neuron comparing single-neuron simulation conditions. Our pipeline can be used to reduce any HH-type multi-compartment neuron model, given its model representation is translated to the NEAT software package. Beyond this, reduction only requires setting of the grid length spacing to obtain adequate spatial sampling of its morphology.

### Validation of spatial reduction performance across varied light conditions and different morphologies

We validated our reduction by comparing the optogenetically induced firing response of the full HH neuron model, serving as a reference (Fig. 2 A), with its response to the somatic-equivalent conductance input, obtained through our reduction (Fig. 2 D, upper panel). This enabled a direct comparison between the full optogenetic simulation and our reduction, as both used the same neuron’s input-output function to transform the input conductance into firing output. Next, to demonstrate the validity of a reduction from the HH to a point-neuron model, we fitted an adaptive exponential integrate-and-fire neuron to the conductance-to-firing curve of the reference HH model (see Methods, *Point-neuron fit to layer 2/3 HH-type model*). When driven by the somatic-equivalent conductance, this point-neuron model reproduced the response of the reference HH model (cf. Fig. 2 D, bottom panel), validating the full reduction.

We next assessed how well the approximation captured the neuron’s complex spatial integration of light. We simulated optogenetic stimulation at different cortical surface locations for the reduced and reference models for a narrow-beam optical fiber (diameter (d)= 50 *µ*m, numerical aperture (NA)= 0.1) at 0.1 mW/mm^2^ stimulation intensity, comparing their spatial response profiles, see Fig. 3 A. The responses of the reduced model matched the reference responses across various stimulator locations, except for a moderate underestimation of the most lateral reference responses. Next, we tested whether the reduced model performed universally well for different light conditions, simulating stimulation with an intermediate, and a divergent fiber (both d = 100 *µ*m, and NA = 0.39 and 0.9, respectively) and varying stimulation intensity (0.13 and 0.25 mW/mm^2^, respectively). Under these varied light conditions, the somatic-equivalent conductance simulated as input to the fitted AdEx point-neuron model well-approximated the reference HH model (cf. 3 A-C, profiles from top to bottom: reference, reduced AdEx, reference minus reduced AdEx).

**Fig 3.**
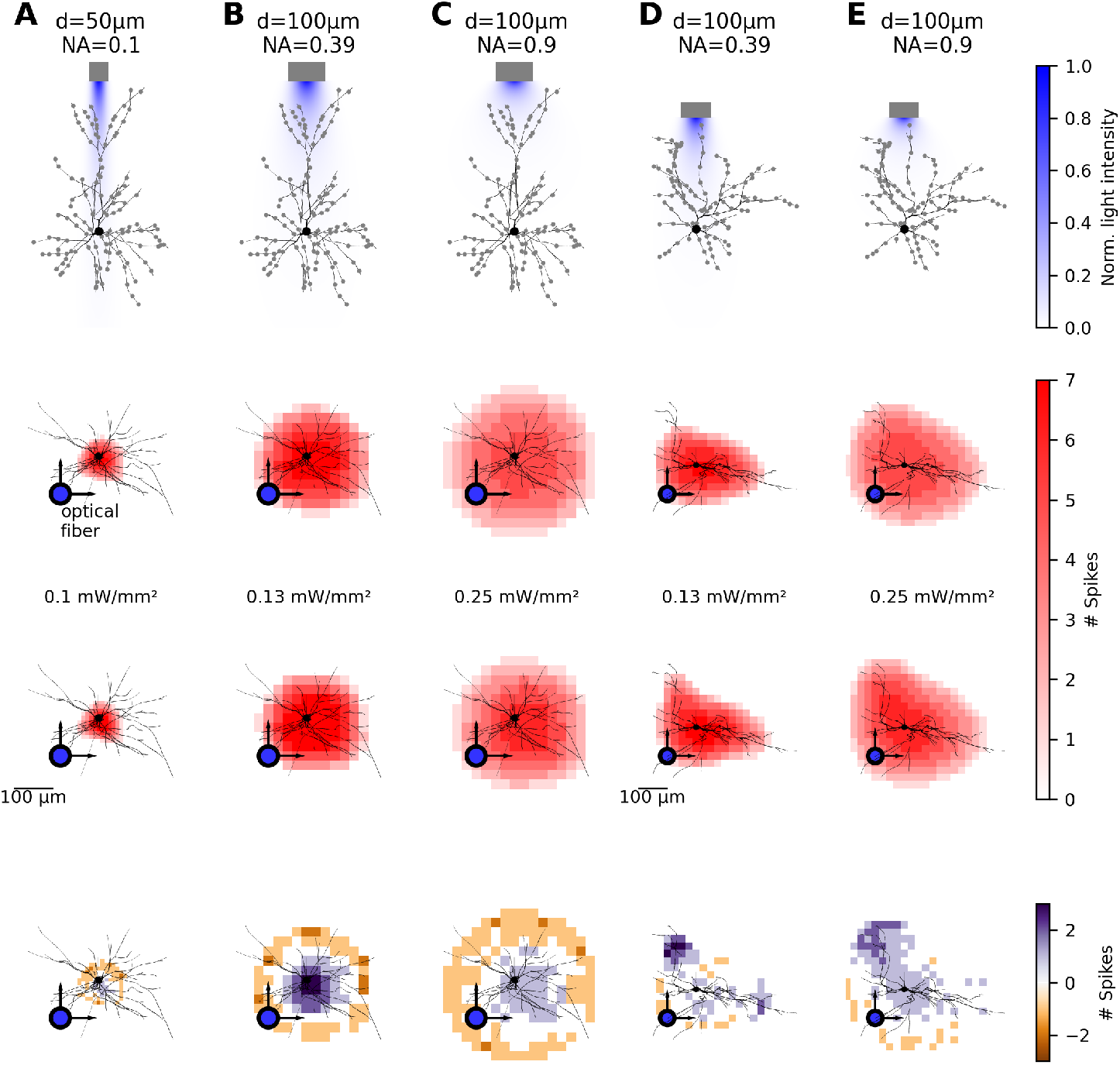
Reduced model predictions of optogenetic responses generalize across light conditions and two neuron morphologies. **(A-E)** Illustration of the simulated optical fiber and reference neuron morphology with its representative spatial reduction as gray dots (top row). Comparison of spatial optogenetic response profiles simulated by varying optical fiber position on top of the cortex for the reference neuron simulations (second row), reduced model simulations (third row), and their difference (reference minus reduced model, bottom row). **(A-C)** Layer 2/3 reference neuron 1 simulated with an optical fiber of low (*d* = 50 *µ*m, *NA* = 0.1), moderate (*d* = 100 *µ*m, *NA* = 0.39), and high beam divergence (*d* = 100 *µ*m, *NA* = 0.9). Stimulation intensity at the fiber output were 0.1, 0.13, and 0.25 mW*/*mm^2^, respectively. Reduced model firing output was simulated using the somatic-equivalent conductance as input to the reference-model-fitted AdEx point-neuron model. **(D,E)** Layer 2/3 reference morphology 2 stimulated by optical fibers with moderate (*d* = 100 *µ*m, *NA* = 0.39), and high beam divergence (*d* = 100 *µ*m, *NA* = 0.9). Stimulation intensity at the fiber output were 0.13 and 0.25 mW*/*mm^2^, respectively. Reduced model firing output was simulated using the somatic-equivalent conductance as input to the reference HH-type model.

Finally, we applied the reduction procedure to another reference morphology of a layer 2/3 pyramidal neuron, while keeping the same grid-width for spatial reduction (50 *µ*m). Simulation of the somatic-equivalent conductance as the only input to the HH-type neuron model matched the results of the full simulations of optogenetic stimulation with this model (cf. see Fig. 3 D,E). These results demonstrate a robust performance of our reduction procedure across two different neuron morphologies and for varied light conditions.

### Morphology sharpens optogenetic activation in V1 network model

Using our reduced models, we simulated optogenetic stimulation of layer 2/3 excitatory neurons in a large-scale model of mammalian primary visual cortex (V1), consisting of cortical layers 2/3 and 4, with feedforward input from model lateral geniculate nucleus (LGN) neurons. The connection probability of model neurons decays laterally according to experimental anatomical observations [19], with local connectivity in layer 4 and long-range connections in layer 2/3. Furthermore, connections follow known functional biases, including push-pull organization in layer 4 and moderate co-orientation bias in layer 2/3. Due to this functionally and anatomically grounded architecture, model neurons express a wide range of realistic functional properties, including spontaneous activity statistics, visual-stimulus orientation tuning, as described by layer-specific predominance of complex and simple cells (cf. Fig. 4 A), and visual-stimulus size tuning [17]. Variations of the model were previously used to study V1 activity under optogenetic stimulation [12, 20], however with the optogenetic mechanism implemented assuming point morphology. Here, we will expand the optogenetic mechanism with the new morphology-aware approximation. To start investigating the impact of optogenetic stimulation on functional cortical networks, we simulated a disk-shaped light-stimulus of varying diameter, projected by a grid of 10 *µ*m-pitch LED stimulation elements positioned on top of model cortex (cf. Fig. 4 B). The optogenetic stimulation targeted only excitatory neurons in layer 2/3, simulating the selective optogenetic transfection of pyramidal neurons.

**Fig 4.**
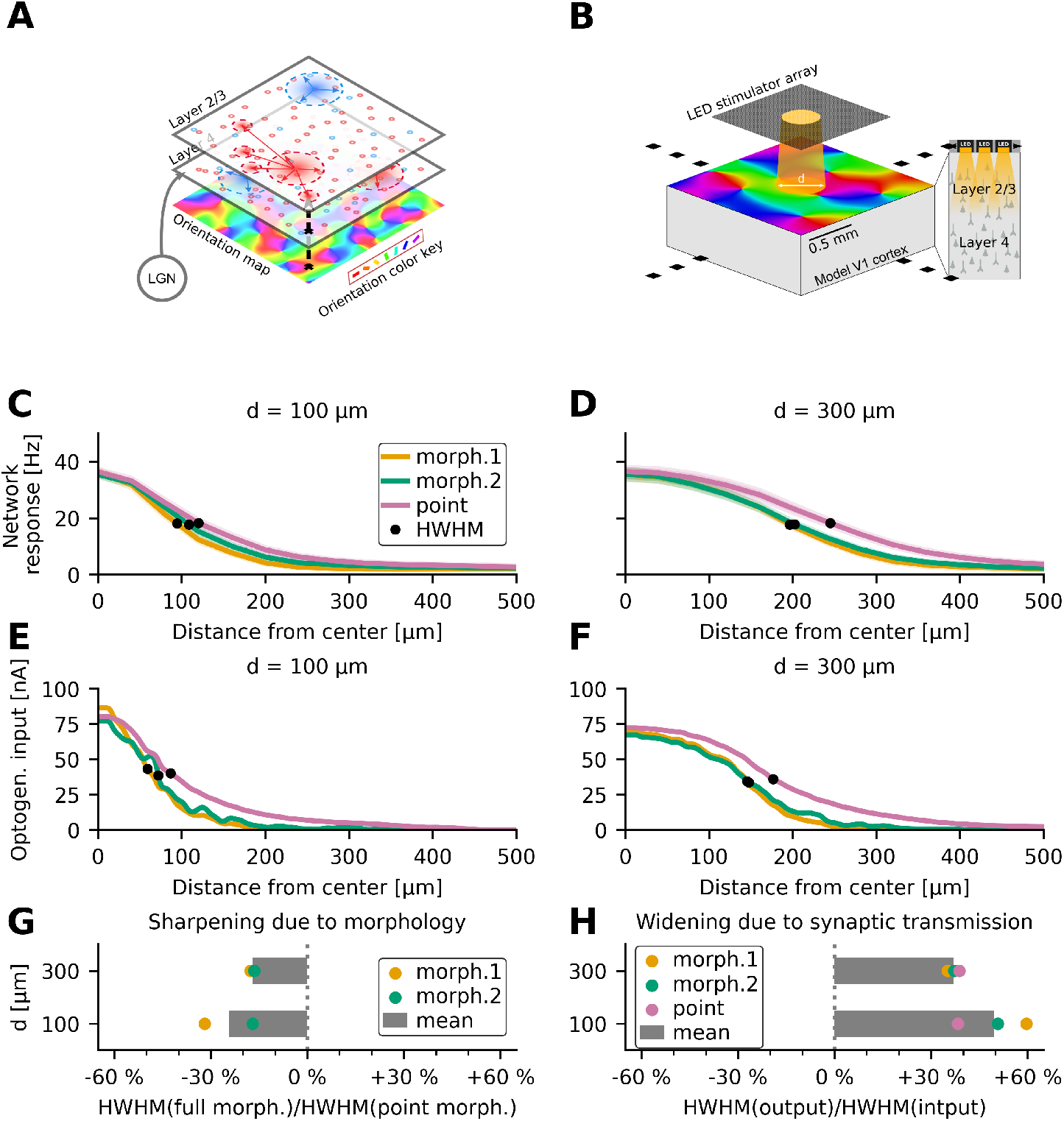
Network mechanisms widen, morphology sharpens optogenetic activation. **(A)** Network model of mammalian primary visual cortex layers 2/3 and 4 built from 93750 adaptive exponential integrate-and-fire neurons, connected based on distance- and neuron-type-dependent connectivity. **(B)** Simulation of light stimulation in model cortex via an LED-stimulator-array (10 *µ*m LED width and spacing). The simulated stimulus is a disk of varying width *d* = [100, 300] *µ*m. Optogenetic stimulation is exclusive to excitatory (pyramidal) neurons in layer 2/3. **(C)** Firing response in the simulated cortical region as a function of distance to center (40 *µ*m resolution), where a 100-*µ*m wide light stimulus is applied. Three conditions were simulated: Optogenetic stimulation of layer 2/3 excitatory neurons with morphology 1 (yellow), morphology 2 (green), and neglecting morphology (“point”, pink). Firing rates were calculated over the stimulation-on interval (100 ms). Their mean (solid curves) and standard error on the mean (SEM, transparent bands) was computed over 30 simulation trials. The half-width-at-half-maximum (HWHM) is marked as a black dot and was evaluated to estimate spatial stimulation resolution. **(D)** Same as (C) for a 300-*µ*m wide light stimulus. **(E,F)** Same as (C,D) for the optogenetic input current into layer 2/3 pyramidal neurons. Simulations were carried out at 2 *µ*m spatial and 1 ms temporal resolution. Since simulations are deterministic, there is no variance across simulation trials. **(G)** Relative sharpening of layer 2/3 neuron’s optogenetic response when simulated with full morphology 1 or 2 (colors) compared to when simulated with point morphology for both stimulus widths (y-axis). Sharpening was calculated by dividing the morphology’s input HWHM by the point morphology’s input HWHM. Grey bars show the mean across both full morphologies. **(H)** Widening attributed to synaptic transmission in the network for realistic and point morphologies (color-coded) as well as both light stimulus width conditions (y-axis). Relative widening is calculated as the ratio of the HWHM of the network response (output) and the HWHM of the optogenetic input. Grey bars show the mean across all morphology conditions.

Surprisingly, across n=30 optogenetic stimulation trials, mean firing rates as a function of distance from the center of the stimulated cortical volume showed that both layer 2/3 pyramidal morphologies produced more spatially confined network activation compared to control simulations with point morphologies, see Fig. 4 C,D. To disentangle whether this paradoxical result was driven by the spatial integration of light by realistic neuron morphologies itself or by a counteracting network mechanism which sharpens evoked network activity, we quantified how the optogenetic input to neurons decayed with distance from the center of the simulated cortical region, see Fig. 4 E,F. We found that optogenetic input to neurons with realistic layer 2/3 pyramidal morphology was spatially more confined than the optogenetic input to neurons with point morphology, indicating that the spatial integration of light was sharper for the former. Evaluation of the half-width-at-half-maximum (HWHM) of the optogenetic input (cf. filled black circles in Fig. 4 E,F), revealed that realistic morphologies sharpened the spatial response profile by 24 % in the narrow (*d* = 100 *µ*m) and by 17 % in the wide input condition (*d* = 300 *µ*m), see Fig. 4 G. Additional simulations, detailed in a later section (*Apical dendrites sharpen optogenetic response spatially*), revealed that this sharpening was caused by apical dendritic contributions, which are driven by the more superficial and thus more spatially confined portion of the light beam.

### Network mechanisms widen spatial spread of optogenetic input by 37-50% depending on input width

To quantify the contribution network mechanisms have in widening optogenetic input, we determined the ratio between the half-width-at-half-maximum (HWHM) of the network response and the optogenetic input (filled black circles in Fig. 4 C-F). Network mechanisms widened the response relative to the optogenetic input by 50 % for narrow input (*d* = 100 *µ*m) and by 37 % for wide input (*d* = 300 *µ*m), see Fig. 4 H. The differences in relative widening were stronger for the narrow input condition (*d* = 100 *µ*m), suggesting a stronger sensitivity of the widening network mechanism to the narrow optogenetic input. In conclusion, synaptic transmission in the network was the only driver of horizontal propagation of optogenetically evoked activity, besides light propagation, in our simulations. Neuronal morphology instead sharpened the evoked activity, partially counteracting the widening effect of the synaptic transmission.

### Small sharpness changes in optogenetic input degrade separation in network output

A key experimental goal in developing cortical visual prostheses is to elicit spatially separable cortical responses to applied stimuli [7, 8]. While the link between cortical activation and perceptual resolution remains under investigation [21], studies using electrical stimulation show that closely spaced inputs evoke fused rather than distinct percepts [22]. This fusion may arise from non-linear network mechanisms that merge responses to nearby stimuli. The minimum distance at which two spatially separate external stimuli evoke spatially separate cortical responses could thus serve as an indicator for the spatial resolution achievable by the visual prosthesis.

We therefore examined how neuron morphology affects responses to two 100 *µ*m-diameter disk-shaped light stimuli placed at varying distances (Fig. 5 A), finding that the evoked response starts to separate at around 200 *µ*m distance. However, at this distance, the firing rate of neurons located between the stimuli remained above 50% of the maximal responses reached right below the two input stimuli, see Fig. 5 C. To quantify separation, we defined the response overlap of both stimuli by the minimum response recorded between them, normalized by the average of their center response. An overlap below 30% was achieved at shorter distances for both pyramidal morphologies (300 *µ*m) than for the point morphology (400 *µ*m), see Fig. 5 C,D. These results indicate that neuronal morphology enhances the separation of responses to paired optogenetic stimuli, consistent with our earlier finding that morphology sharpens responses to single stimuli. More generally, this highlights that even moderate improvements in precision of paired inputs, whether due to morphology or other factors, can critically enhance the separation of network outputs.

**Fig 5.**
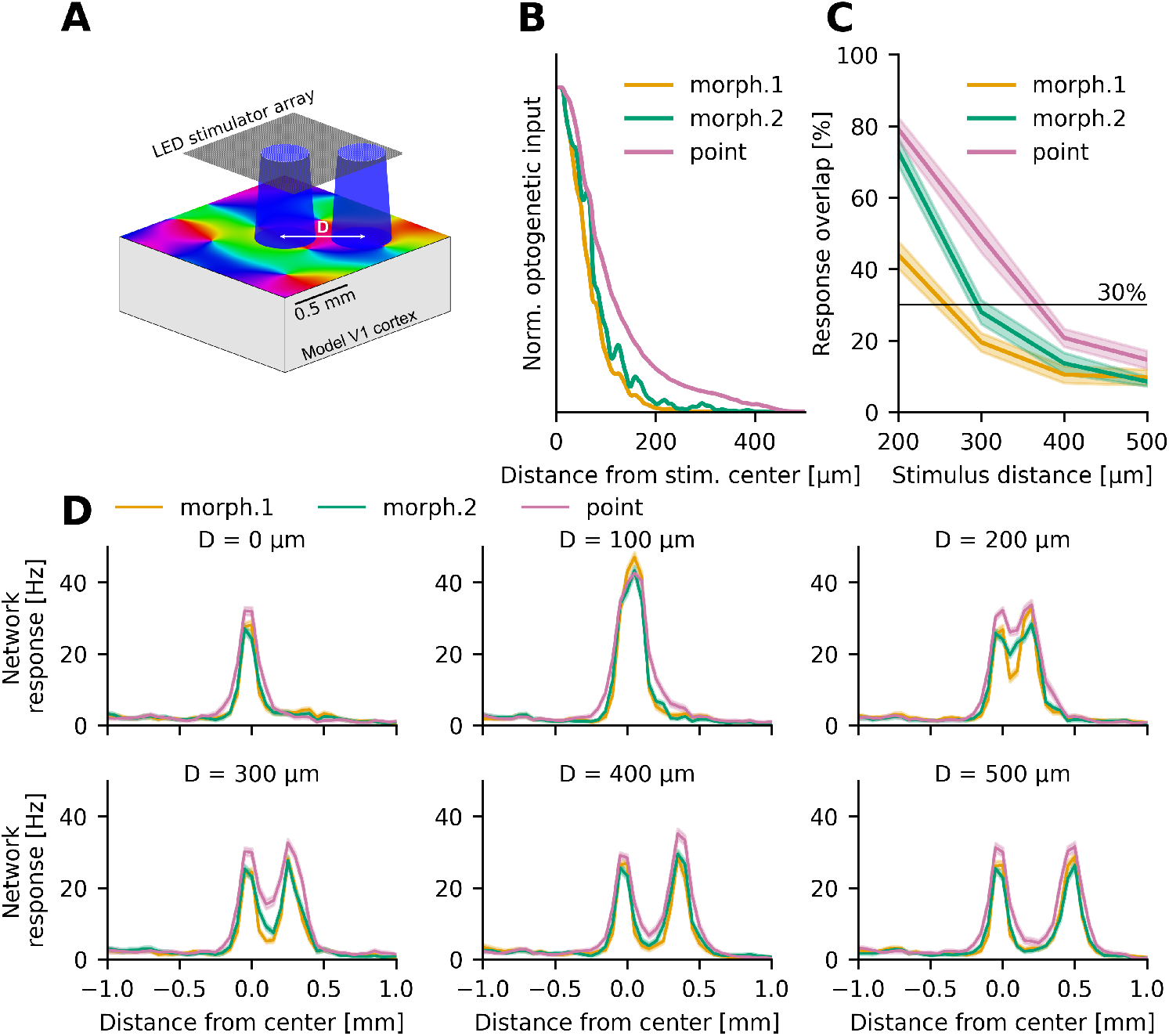
Moderate widening of network input can break separation of two spatially separated input stimuli in network output. **(A)** Two 100 *µ*m-wide input-stimuli separated by a distance *D* between their centers were applied to the model V1 cortex. Stimulation intensity was adjusted to evoke 30-35 Hz firing response at stimulus center. **(B)** Normalized optogenetic input plotted over distance from stimulus center for both morphology and the point-neuron conditions. **(C)** Overlap of the two individual optogenetic stimuli in network output as function of the distance between stimuli centers. We define overlap by the network output at the minimum located between the two stimuli centers normalized by the average network output at the two stimuli centers. **(D)**. One-dimensional profiles of the network output for a single stimulus (*D* = 0*µ*m) and for two simultaneously presented input stimuli with center distances of *D* = 100 - 500 *µ*m in steps of 100 *µ*m measured along the line spanned by the stimuli centers.

### Apical dendrites sharpen optogenetic response spatially

To disentangle why realistic layer 2/3 neuron morphology caused sharper optogenetic responses than point morphology, we compared the somatic-equivalent conductance contributed by different morphological parts. We hypothesized that the sharpening was driven by the apical dendrite, which, due to its superficial location, is exposed to a more confined light beam than the basal dendrites or soma, where the light beam widens due to dispersion. Hence, when apical contributions make up a substantial portion of the total optogenetic input, the full pyramidal morphology receives a sharper input than the point morphology. We tested this hypothesis by splitting the morphology into apical and basal parts at 100 *µ*m distance above soma (cf. Fig. 6 A). We then simulated the somatic-equivalent conductance for the full, apical-only, and basal-only morphology, as well as for the point morphology. Using an optical fiber with wide beam (*d* = 100 *µ*m, *NA* = 0.9) mimicking the divergent light emission profile of the LED array employed in the network simulations, we found that the total (time-integrated) apical contribution decayed faster with light-source distance than the basal and somatic contributions, see Fig. 6 B. Repeating the simulations for the second morphology confirmed these observations (cf. Fig. 6 C, D). Consistent with our hypothesis, apical dendritic contributions improved the sharpness of the total optogenetic input to the realistic morphology compared to the point morphology, see Fig. 6 B, C, ‘apical’, ‘full’ vs. ‘point’. Given that stronger confinement of the light beam at apical versus basal dendritic depth caused the precision advantage, we repeated simulations with a less divergent optical fiber (*d* = 100 *µ*m, *NA* = 0.39), confirming that the sharpening effect vanishes, see Fig. 6 E, F. Overall, these results confirm that for stimulation with divergent light beams, apical morphology has a sharpening impact on a layer 2/3 pyramidal cell’s optogenetic input.

**Fig 6.**
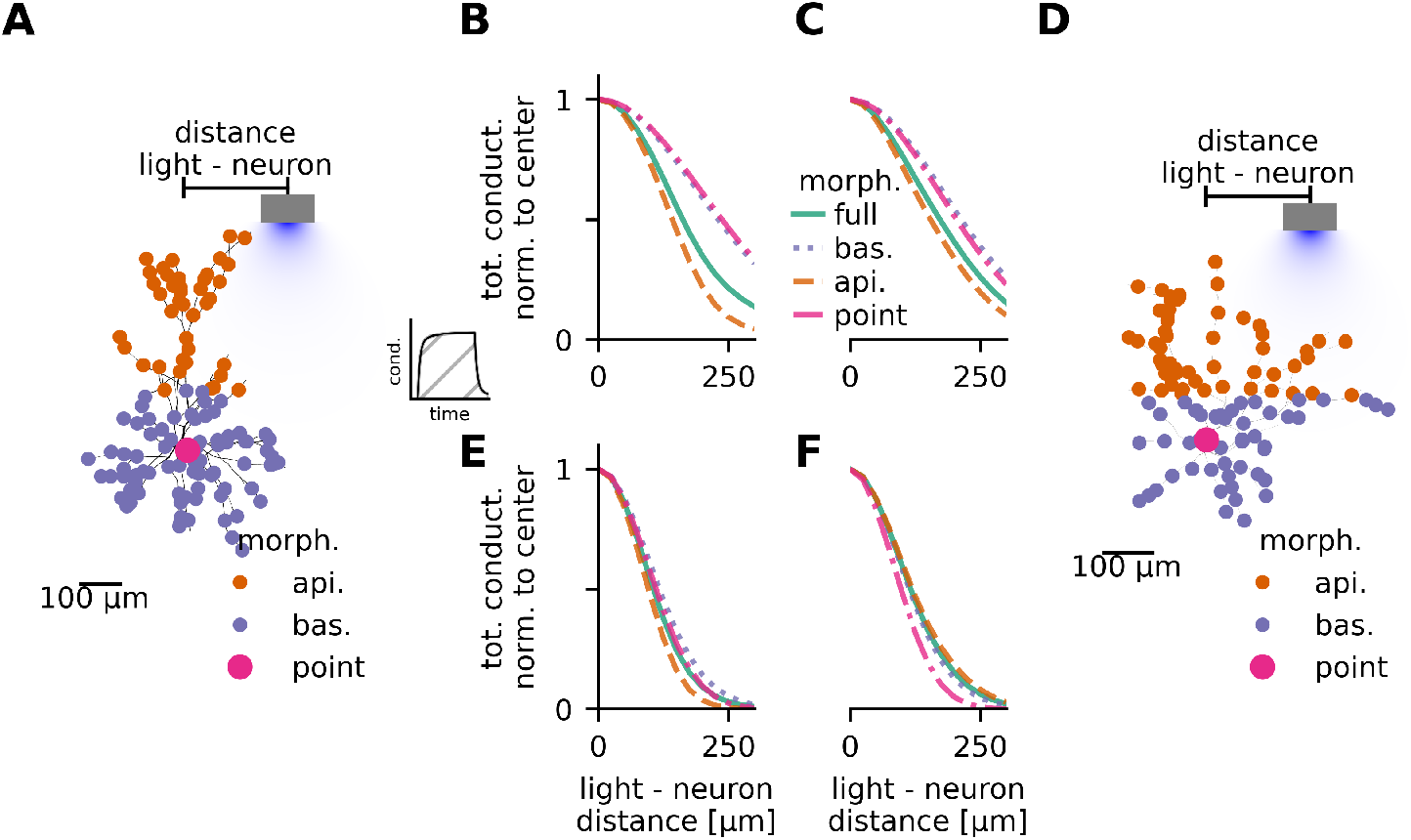
Apical dendrites sharpen the optogenetic response of layer 2/3 pyramidal neurons. **(A)** Optogenetic stimulation of layer 2/3 pyramidal neuron model 1 with a wide optical fiber (*d* = 100 *µ*m, *NA* = 0.9). Somatic-equivalent optogenetic conductance was calculated separately for apical, basal, and somatic compartments (color-coded). **(B)** Normalized and time-integrated somatic-equivalent conductance (see inset) depending on radial distance between neuron and fiber (x-axis). Conductance origin (apical, basal, somatic, full morphology) color-coded like in (A). **(C,D)** Same as (B) and (A) for layer 2/3 neuron model 2. **(E,F)** Repetition of simulations from (B,C) for narrower optical fiber with *d* = 100 *µ*m and *NA* = 0.39. Stimulation intensity was 0.2 mW/mm at optical fiber output for all conditions.

## Discussion

Optogenetic stimulation enables the study of information processing in brain circuits [3, 4, 11] and could open a path for restoring sensory function through next-generation cortical prosthetics [10, 12]. To engage with spatially encoded sensory code in the cortex, optogenetic techniques must cover a wide field of cortex while enabling precise control over small spatial volumes [7, 1]. Spread of evoked activity beyond the targeted volume presents a crucial limitation for such precise control [3], which an improved mechanistic understanding could help to resolve. To address this, we introduced a comprehensive model of cortical optogenetic stimulation in primary visual cortex, which accounts for the light dispersion in cortical tissue, neuron morphology, and network mechanisms, jointly in a single model.

Our simulations predict 37-50% widening of evoked activity compared to initial stimulus width, which is in good agreement with experimental results from tree shrew V1, observing a 34-80% wider extent of evoked activity in this species [3]. Our predictions are lower than those of a previous computational point-neuron-based network model simulating a two-dimensional neuronal sheet with uniform connectivity, finding 50-150% widening [16]. Notably, our model suggests that, besides light scattering, only synaptic transmission of the optogenetically evoked activation in the network drives spatial widening of evoked activity, whereas, paradoxically, spatial activation of layer 2/3 pyramidal neuron morphology has narrowing impact. Our single-neuron simulations revealed that it is the apical dendrites sharpening the optogenetic input to the cell, over-compensating the broader input received by the basal dendrites and soma. This single-neuron effect explained why neuron models using realistic morphology predicted narrower activation than models using point-morphology in our network simulations.

Our study offers valuable insights for future experiments featuring divergent light stimulation. First, optogenetic input to the entire morphology of layer 2/3 pyramidal cells was more precise than optogenetic input to their soma alone. This suggests that somatic targeting of channelrhodopsin is unlikely to improve stimulation precision, while it requires higher stimulation intensity [13], increasing the risk of phototoxicity. Instead of somatic targeting, future studies could try optimizing light profiles to preferentially target apical over basal dendrites, enhancing their sharpening effect. Second, network effects contributed to spatial spread via non-linear input-dependent mechanisms. When a single light stimulus was presented, network effects widened initially narrow stimuli relatively more than wider stimuli. However, when two spatially separated input stimuli were presented, their separation in network output improved significantly with narrower initial input width. Hence, delivering of narrow input stimuli could be more critical for spatial separation of simultaneously applied stimuli while being less crucial for inducing confined activation with a single stimulus.

Our work also tested whether simulations of optogenetic stimulation in cortical networks require modeling of layer 2/3 neuron morphology explicitly to deliver accurate results. Our results suggest that point-neuron models enable capturing core network effects which are the dominant determinant of the activation spread beyond the illuminated volume, however, morphology becomes relevant for finer-scale effects, such as under optogenetic stimulation with complex light patterns.

Future studies can build on our morphology approximations or use our procedure to derive their own to integrate morphology-aware optogenetic stimulation into existing point-neuron-based network simulations. To that end, the software pipeline implementing the reduction procedure and the reduced models of two layer 2/3 pyramidal neurons are published along this article. Simulation of reduced models requires only a minimal Python software installation (Python and Numpy), enabling a smooth integration into any Python-based network simulation. The channelrhodopsin model and distribution inside the neuron’s morphology can be adapted to simulate specific experiment scenarios. Finally, the reduction procedure can be generalized to build reduced models of other neuron types as it is designed in a straight-forward manner, only requiring adjusting the grid-length for spatial reduction of the original morphology. Additionally, the full ready-to-use implementation of the reduced models within the network simulations is made freely available as part of the Mozaik package [23].

The main limitation of this study is that we approximate dendritic-filtering neglecting inter-compartment interactions, which introduces prediction errors on how strongly various dendritic parts contribute to activation. While we demonstrate that a global scale factor compensates the prediction error for layer 2/3 pyramidal cells, approximation precision could be reduced for reductions of larger neurons like layer-5 pyramidal cells as errors increase with distance between compartmental sites. However, imprecision due to large dendritic distances could be resolved through computing interactions as outlined in Wybo et al. at the cost of increased computation time.

## Methods

### Morphological L2/3 pyramidal neuron models

Multi-compartment neuron models of two layer 2/3 pyramidal neurons of rat somatosensory cortex were obtained from Markram et al. [24] and Ramaswamy et al. [25] (identifiers: “L23 PC cADpyr229 1” and “L23 PC cADpyr229 3”, see https://bbp.epfl.ch/nmc-portal/microcircuit.html#/metype/L23_PC_cADpyr/details for details). Neurons were implemented in NEAT [18] (https://neatdend.readthedocs.io/en/latest/index.html) and simulations carried out with NEURON [26] (https://www.neuron.yale.edu/neuron/).

### Optogenetic conductance model

We modeled conductance dynamics of ChrimsonR as captured in a 5-state Markov model. This model was fitted to patch-clamp experiments performed on ChrimsonR-expressing HEK293 cells [27]. The model assumes two open (O1, O2), two closed (C1, C2), and one slow state (S). Light induces molecule transition from closed to open states, see Fig. 7 and affects switching between closed states. The molecule relaxes back from open to closed states and further from open state 2 to closed state 1 via the slow state at light-independent rates. The ratio of conductance in the open states was 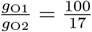 [27].

**Fig 7.**
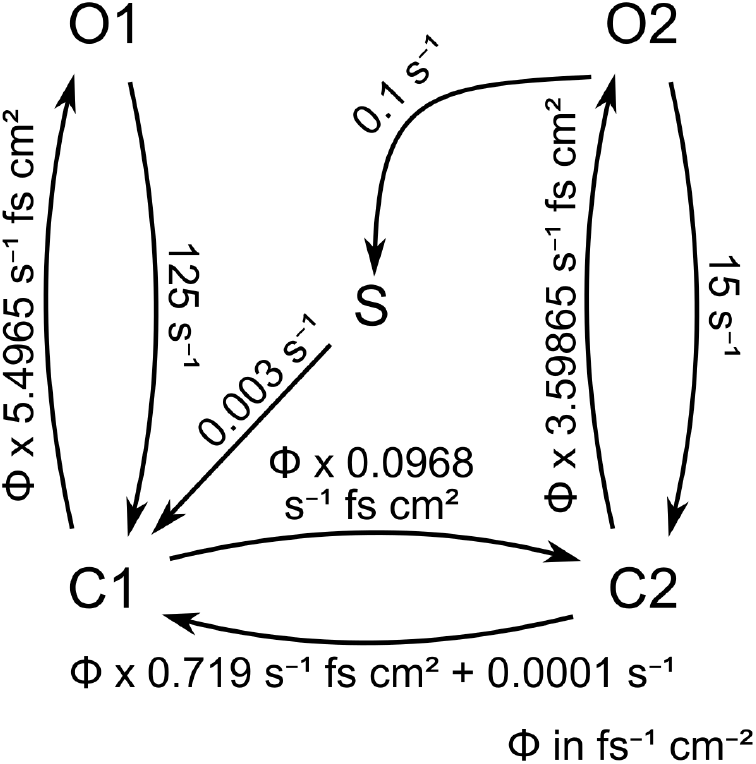
5-state Markov model of ChrimsonR molecule conformation changes under light exposure [27]. The model consists of closed states, C1 and C2, open states, O1 and O2, as well as a slow state, S. Light-dependent switching rates have a linear dependency on photon fluxΦ.

For full and reduced simulations of layer 2/3 pyramidal cells, ChrimsonR was uniformly expressed at 130 channels*/µ*m^2^ across the neurons. Single-channel conductance in O1 was 50 fs based on measurements of bacteriorhodopsin [28]. Conductance in O2 was scaled to match the ratio of conductance between the open states to 8.5 fs. In the network simulations, we simulated neurons with a ChrimsonR expression of 15 channels*/µ*m^2^ in the full morphology conditions, approximately matching the total conductance in open states 1 and 2 of the point morphology condition (17 ns and 2.9 ns, respectively).

### Morphology reduction

#### Somatic-equivalent optogenetic conductance

To approximate the effect of morphology on a neuron’s optogenetic response without simulating the entire morphology with a multi-compartment approach, we derive the equivalent optogenetic somatic conductance. This represents the aggregate somatic influence of conductance distributed over the entire dendritic tree. Hence, we need to find a rescale factor that approximates dendritic attenuation in order to relocate a dendritic conductance to the soma or into a point neuron model instead.

To enable the analytic derivation of transfer-impedance across the neuron’s morphology, we absorb the conductance at equilibrium of all active dendritic ion channels into the passive leak [18]. The impact of a current *i* at a dendritic site *d* on the membrane voltage *v*_*s*_ at the soma can be expressed as the convolution of the time-dependent transfer-impedance between dendrite and soma with the input current:

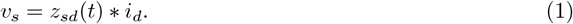

Wybo et al. [18] have shown that the temporal evolution of the dendro-somatic transfer impedance can be approximated by the temporal evolution of the somatic input impedance. As the latter is applied implicitly during simulation, a rescale factor can be used to correct for the difference in driving force between soma and dendrite. Specifically, an optogenetic conductance, *g*(*t*), introduces a current *i*(*t*) in a dendritic site depending on the site’s membrane voltage *v*_*d*_(*t*) and the conductance’s reversal potential *e*. The impact of this single conductance at soma can be described by:

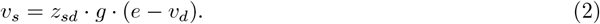

To relocate the conductance to soma, a rescale factor *β* has to be found such that the dependence on local dendritic voltage is replaced with a somatic voltage dependence:

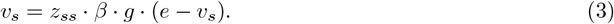

Under the aforementioned assumption that the transfer impedance between dendritic site and soma can be approximated by the somatic input impedance (*z*_*sd*_ ≈*z*_*ss*_), which is justifiable for neurons with small apical dendrites, such as pyramidal neurons of layer 2/3 [29], the rescale factor becomes [18]:

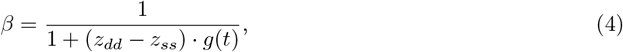

with the dendritic input impedance *z*_*dd*_.

Note that this derivation assumes that input is present at one dendritic site only. Simultaneous input at another site can increase the membrane voltage at the input site, diminishing its actual impact at soma. Generalizing the rescale factor to parallel input at various sites requires solving a system of linear equations of the dimension equaling the number of compartments (≥1000) for each time step, see Wybo et al. [18], which in turn violated our computation performance requirements. We therefore approximated the somatic-equivalent conductance neglecting the coupling between dendritic compartments and compensated the overestimation of dendritic impact at soma through a global scale factor,

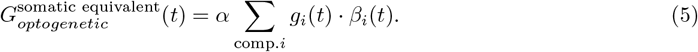

To determine the optimal value of the scale factor *α*, we compared the somatic-equivalent conductance calculated using our approximation and the full calculation from Wybo et al. [18], finding that a constant factor of *α* = 0.85 robustly captured neglected inter-compartment couplings (see Fig. 2 C).

#### Spatial reduction of neuron morphology and approximation of somatic equivalent conductance

To speed up model computation, we reduced the number of morphology locations at which the light intensity and respective optogenetic conductance are computed. To ensure representative sampling of the entire neuron, we aimed to balance two goals: (1) even coverage along the morphological structure and (2) inclusion of spatially distal segments that would be missed by sampling at fixed steps along the morphology’s trajectory alone. We achieved this by selecting morphology points closest to nodes of a Cartesian grid with 50 *µ*m spacing. This approach prevents oversampling of densely clustered regions and ensures that sparsely located, distal branches are also captured.

To preserve the reference neuron’s full electrophysiological properties, we calculated the optogenetic conductance in all compartments of the reference neuron based on their membrane area and local channelrhodopsin expression, but used the single-channel conductance at the nearest sampled location instead of calculating this conductance for each compartment individually. The final somatic equivalent conductance was then computed by rescaling of and summing over these values, as described in eq. 5.

### Point-neuron fit to layer 2/3 HH-type model

We adopted the adaptive exponential (AdEx) integrate-and-fire neuron model [30] to reproduce the input-output curve of the full HH-type layer 2/3 pyramidal cell (reference morphology 1, model identifier: “L23 PC cADpyr229 1”) under exposure to 200 ms conductance profiles of varying strength (cf. Fig. 8 A). A multidimensional parameter sweep varying membrane characteristics (capacitance, leak, refractory period) and the spike-triggered adaptation threshold was conducted to identify the most suitable model configuration, see Table 1. The spike response of the fitted AdEx point neuron and the original HH-type neuron are compared in Fig. 8 B.

**Table 1.** Parameters of the adaptive exponential integrate-and-fire neuron model fitted to reference morphology 1.

| Parameter | Description | Value |
| --- | --- | --- |
| $C_m$ (pF) | Membrane capacitance | 90 |
| $\tau_{\text{ref}}$ (ms) | Refractory period | 7 |
| $V_{\text{reset}}$ (mV) | Reset potential | −60 |
| $g_L$ (nS) | Leak conductance | 2.4 |
| $E_L$ (mV) | Leak reversal | −80 |
| $a$ (nS) | Subthreshold adaptation | −0.18 |
| $b$ (pA) | Spike-triggered adaptation | 300 |
| $\Delta_T$ (mV) | Slope factor | 0.8 |
| $\tau_w$ (ms) | Adaptation time constant | 10 |
| $V_{\text{th}}$ (mV) | Threshold potential | −55.5 |
| $V_{\text{peak}}$ (mV) | Spike detection threshold | −40 |

**Fig 8.**
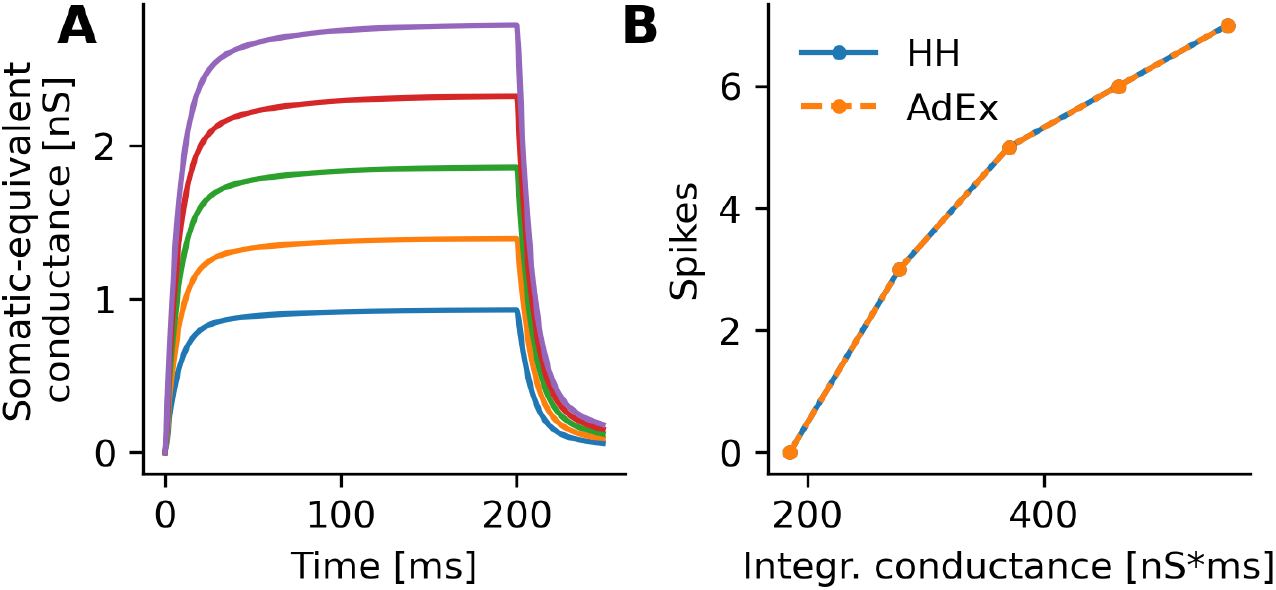
Fitting the adaptive-exponential integrate-and-fire (AdEx) model to the reference HH-type layer 2/3 pyramidal neuron 1. **(A)** Conductance input profiles of 200 ms-duration with varying strength. **(B)** Spiking responses of the fitted AdEx point-neuron model (orange) and the original HH-type model to the inputs in (A) plotted against the integral of the input conductance.

### Optical fiber light model

We simulated light emission from an optical fiber in cortical tissue using the Kubelka-Munk theory, accounting for absorption (0.125 mm^−1^) and scattering (7.37 mm^−1^) [13, 14, 31], with a refractive index of 1.36 [32]. We assumed direct placement of the fiber on top of cortex and no coupling losses. Fiber diameter and numerical aperture are model parameters, which define the output surface and beam divergence.

### Single-neuron simulations with full and reduced morphology

Optogenetic responses of layer 2/3 pyramidal neurons were simulated for both reference multi-compartment Hodgkin-Huxley(HH)-type neurons (described in ‘L2/3 pyramidal neuron models’) and using their counterparts with reduced morphology, developed following the procedure described in ‘Morphology reduction’. For simulations of neurons with reduced morphology, the somatic-equivalent conductance was simulated as the only conductance input to either the reference HH-type neuron or to a point-neuron model, which had been fitted to the reference HH-type neuron beforehand.

To simulate spatial response profiles, cell morphologies were placed at appropriate cortical depths (cell 1: 423 *µ*m, cell 2: 389 *µ*m). Responses to optogenetic stimulation applied from the cortical surface were simulated with an optical fiber light emission profile (see ‘Optical fiber light model’) and 200 ms-duration constant illumination. Simulations were repeated for varying stimulator positions defined by a two-dimensional grid on the cortical surface. The grid spanned *xy* = [−110, 100], *dxy* = 10 for simulations presented in Fig. 2 and *xy* = [−300, 275], *dxy* = 25 in Fig. 3.

### Network simulations

#### Mammalian primary visual cortex network model

To explore how the simulation of realistic versus point morphology affected network-level optogenetic stimulation responses, we used an anatomically and functionally constrained large-scale model of mammalian primary visual cortex (V1) [17]. Here, we describe the core architecture of the model and refer the reader to the original publication for implementation details [17].

The model consists of six populations of adaptive exponential integrate and fire neurons: lateral geniculate nucleus (LGN) ON and OFF cells, and cortical excitatory and inhibitory layers 4 and 2/3. LGN cells receive current input corresponding to the convolution of visual input with their spatiotemporal receptive fields, scaled by contrast and luminance. These LGN cells provide feedforward input to layer 4, with connectivity based on receptive field templates assigned to layer 4 cells based on a pre-computed orientation preference map [33]. Intra-layer 4, excitatory neurons preferentially connect to excitatory neurons with correlated receptive fields and inhibitory neurons with anticorrelated receptive fields, following push-pull [34] organization. Layer 2/3 neurons receive inputs from layer 4 neurons, and provide feedback to them, as substitution for the biological feedback path through infragranular layers 5 and 6, which are not included in our model. In addition, layer 2/3 neurons form long-range intra-layer connections biased towards iso-oriented neurons. Intracortical connectivity is constrained by experimentally measured connection probability density profiles [35, 36]. For additional details, see Antolík et al. 2024 [17].

#### Optogenetic stimulation framework for the network model

We simulated optogenetic stimulation in the network conditions using a computational model of a µLED-based optogenetic stimulator [12], which we expanded by the reduced models of layer 2/3 morphology. The simulated LED array was positioned above the cortical surface, covering 4 × 4 mm^2^ of its area, with directly neighboring LED elements of 10 *µ*m diameter. Light propagation in cortical tissue was simulated for light of 590 nm wavelength using the ‘Human Brain Grey Matter’ model implemented in the LightTools software. Light scattering was modeled with the Henyey-Greenstein model [37] using *g* = 0.87 as anisotropy factor and a mean free path of 0.07 mm [38].

### Statistics and data analysis

#### Network simulation responses and optogenetic input currents

To quantify the network response across morphologies and optogenetic stimulus radii, we recorded model spikes on a 4×4 mm grid of electrodes with 40 µm spacing. For each electrode, we averaged spike counts of neurons in a 50 µm radius every 5 ms, resulting in the mean firing rate of neurons in the local vicinity. We applied optogenetic stimulation within circular regions of 100 µm and 300 µm diameter, for 100 ms with a 350 ms inter-stimulus interval and 30 trials centered on the middle of the layer 2/3 cortical sheet. We subsequently calculated the radial mean around this center point and temporal mean for each stimulation radius. For each stimulation radius we calculated the standard error of the mean of the resulting signal across trials.

To quantify the spatial extent of the total optogenetic input current injected into our model cortex neurons (Fig. 4 E,F), we retrieved the inserted current time course and cortical position of each stimulated neuron. We summed the current across stimulation time points, and calculated the distance of each neuron from the stimulation center. To acquire a continuous relationship of radial distance and injected current from these data points, we applied Gaussian smoothing with a standard deviation of 5 µm along the radial distance dimension. The input current was the same for each neuron for all trials with the same stimulation radius.

#### Response overlap of two simultaneous optogenetic responses

We simultaneously applied two disk-shaped optogenetic stimuli of 100 µm-diameter for 100 ms with a 350 ms inter-stimulus interval and 30 trials. One disk was centered on the middle of the layer 2/3 cortical sheet and another one at varying center-distances *D* = [0, 500] *µ*m, Δ*D* = 100 *µ*m). We recorded model spikes as outlined above for the previous network simulation experiments and evaluated mean and standard deviation over simulation trials.

To quantify the overlap of the mean network responses to the two simultaneously applied stimuli, we evaluated the response at the minimum between the stimuli and normalized by the average of the two peak responses, being observed centrally below the two stimuli. Standard deviations of response overlap were obtained by propagating trial-to-trial variability in the network response.

#### Analysis of apical, basal, and somatic contributions to optogenetic conductance

We simulated somatic-equivalent contributions of apical, basal, and somatic parts of the reduced layer 2/3 pyramidal neurons for a 200 ms constant optogenetic stimulus applied through a 100 *µ*m-diameter optical fiber with numerical aperture of 0.9 and 0.39 for the wide and narrow beam conditions, respectively. The fiber was placed on top of the cortical surface at radial distances of 0 - 300 *µ*m in steps of 25 *µ*m and in 20 equidistant angles spanning 2*π* relative to the neuron morphology. We calculated the total conductance in apical, basal, somatic parts and the full morphology by integrating over time. We averaged the total conductance of each morphology part across simulated angles to obtain the average decay of conductance with increasing distance between neuron and light source. We subsequently normalized the decay by the total conductance observed at the central light source position, to obtain the relative decay of each contribution with distance.

## Data and software availability

The morphology reduction and simulation software is implemented in Python and available at (LINK). The network model of primary visual cortex and its optogenetic stimulation setup, which contains the simulation of reduced morphologies as a built-in feature, are implemented in the Mozaik framework [23], which is freely accessible at (LINK). Source code for morphology reduction, single-neuron and network simulations conducted for this study is available at (LINK).

## Acknowledgments

We thank Joshua Böttcher for the translation of neuron model implementations into NEAT.

